# Classic and dissociative psychedelics induce similar hyper-synchronous states in the cognitive-limbic cortex-basal ganglia system

**DOI:** 10.1101/2022.09.27.509527

**Authors:** Ivani Brys, Sebastian Barrientos, Jon Ezra Ward, Jonathan Wallander, Per Petersson, Pär Halje

## Abstract

The neurophysiological mechanisms behind the profound changes in perception and cognition induced by psychedelic drugs are not well understood. To identify neuronal activity specific to the psychedelic state, we here investigated the effects of classic psychedelics (LSD, DOI) and dissociative psychedelics (ketamine, PCP) on neuronal firing rates and local field potentials in several brain structures involved in cognitive processing in freely moving rats.

The classic psychedelics had a net inhibitory effect on firing rates of putative interneurons and principal cells in all recorded regions. The dissociative psychedelics had a similar inhibitory effect on principal cells, but an opposite excitatory effect on interneurons in most regions. However, the inhibitory effect on principal cells was not specific to the psychedelic state, as similar inhibition occurred with a non-psychedelic psychotropic control (amphetamine).

In contrast, both types of psychedelics dramatically increased the prevalence of high-frequency oscillations (HFOs) in local field potentials, while the non-psychedelic control did not. Further analysis revealed strong HFO phase locking between structures and very small phase differences corresponding to <1 ms delays. Such standing-wave behavior suggests local generation of HFOs in multiple regions and weak, fast coupling between structures.

The observed HFO hypersynchrony is likely to have major effects on processes that rely on integration of information across neuronal systems, and it might be an important mechanism behind the changes in perception and cognition during psychedelic drug use. Potentially, similar mechanisms could induce hallucinations and delusions in psychotic disorders and would constitute promising targets for new antipsychotic treatments.

## INTRODUCTION

Converging evidence show that psychedelics are effective at treating several neuropsychiatric conditions and that their therapeutic effect depends mainly on their ability to induce neuroplasticity (Nutt et al., 2020; Vargas et al., 2021; Vollenweider & Preller, 2020). Less is known about how psychedelics alter brain activity to induce the acute changes in perception and cognition that they are most known for. The acute psychedelic state is important to study in a medical context, both for its potential contribution to long-term therapeutic effects but also as a model for psychosis. More fundamentally, psychedelics-induced changes in brain activity might reveal processes important for the study of consciousness (Yaden et al., 2021).

Psychedelic drugs are primarily classified phenomenologically based on their ability to induce a psychedelic experience. Nevertheless, it is well established that classic psychedelics like lysergic acid diethylamide (LSD) and 2,5-dimethoxy-4-iodo-amphetamine (DOI) exert their effects mainly through their agonistic action on 5-HT2A receptors (González-Maeso et al., 2007; López-Giménez & González-Maeso, 2017; Vollenweider et al., 1998), while dissociative psychedelics, such as ketamine and phencyclidine (PCP), act mainly as non-competitive antagonists on N-methyl-D-aspartate (NMDA) glutamate receptors (Jentsch & Roth, 1999; Li & Vlisides, 2016). Despite these differences, increased glutamatergic neurotransmission in corticolimbic networks has been identified as a common downstream effect linked to the psychedelic state (de Gregorio et al., 2018; Nichols, 2016; Vollenweider & Kometer, 2010). In particular, the psychedelic state has been linked to glutamate-dependent depolarizing membrane currents in a subpopulation of pyramidal cells in medial prefrontal cortex (mPFC; Aghajanian & Marek, 1997; Béïque et al., 2007; González-Maeso et al., 2007). However, this increased glutamate signaling does not result in general network excitation and the effect on neuronal spiking activity is complex. The very few investigations performed in awake animals to date indicate that the highly selective 5-HT2A agonist DOI decreases spiking activity in the orbitofrontal cortex, anterior cingulate cortex and motor cortex of rodents (Rangel-Barajas et al., 2017; Wood et al., 2012). In contrast, the NMDA antagonists ketamine, PCP and MK801 promote net excitation in rat mPFC, even if many individual cells also respond with inhibition (Amat-Foraster et al., 2019; Furth et al., 2017; Homayoun & Moghaddam, 2007; Jackson et al., 2004; Suzuki et al., 2002; Wood et al., 2012).

Investigations of synchronized neuronal activity, in the form of local field potentials, have more consistently found overlapping effects for both classic and dissociative psychedelics. Ketamine has been found to induce aberrant high-frequency oscillations (HFO, 110 – 180 Hz) in several corticolimbic structures in rodents (Hakami et al., 2009; Hunt et al., 2006, 2011; Kulikova et al., 2012; Nicolás et al., 2011). Similarly, 5-HT2A agonists induce HFOs in the ventral striatum and mPFC of rodents (Goda et al., 2013; Hansen et al., 2019). HFOs occur to a lesser degree in normal conditions in some of these structures and are believed to enable the integration of otherwise isolated neuronal information through synchronous activity (Buzsáki & Silva, 2012; Eckhorn et al., 1988; Sompolinsky et al., 1990). The olfactory bulb has been identified as an important source of psychedelic-induced HFOs (Hunt et al., 2019; Wróbel et al., 2020). However, local infusion experiments suggest that HFO generation can be initiated independently in multiple structures and then spread to other regions (Hunt et al., 2010; Lee et al., 2017).

Here, we investigated spike activity and HFO oscillations in recordings from several brain structures in parallel in freely behaving rats to identify both local and system-wide neuronal activity changes specific to the psychedelic state. We compared classic psychedelics (LSD, DOI) and dissociative psychedelics (ketamine, PCP) to a non-psychedelic psychoactive control (amphetamine) and found that aberrant HFOs were consistently and specifically present during the psychedelic state in ventral striatum and in medial prefrontal, orbital and olfactory cortices. Moreover, we found that HFOs were phase synchronized between brain structures in a way that could severely influence the integration of information across neuronal systems. Hence, we propose that such hypersynchrony is a mechanism by which an altered state of consciousness can be induced, either during a psychedelic experience or as part of a psychotic episode.

## MATERIALS AND METHODS

### Animals

Nine Sprague-Dawley rats were used (Taconic, Denmark) in this study. All procedures were approved in advance by the Malmö/Lund ethical committee of animal experiments.

### Construcion and Implantation of electrode arrays

Microelectrode arrays with 128 wires were built and implanted as previously described (Ivica et al., 2014). Three screws on the occipital bone (over the cerebellum) were connected to a silver wire and used as ground for the recording system. The animals were allowed to recover for at least two weeks ater implantation.

Electrode positions were verified post mortem in 5 animals using computed tomography (see Supplementary methods). A total of 50 unique anatomical labels were attributed to the electrode tips in this way and they were further grouped into 10 broader regions based on assumed functional similarity (see Table S1 and Table S2).

### Pharmacological treatments

To record the behavioral and electrophysiological effects of pharmacological treatments, animals were placed in a round open field arena. After ∼60 min of baseline recording, the animal was intraperitoneally injected with LSD (lysergic acid diethylamid, 0.3 mg/kg, Lipomed AG, Switzerland), DOI (2,5-dimethoxy-4-iodoamphetamine hydrochloride, 2 mg/kg, Lipomed AB, Switzerland), ketamine (Ketaminol, 25 -50 mg/kg, Intervet AB, Sweden), PCP (phencyclidine hydrochloride, 5 mg/kg, Lipomed AG, Switzerland) or amphetamine (d-amphetamine hydrochloride, 4 mg/kg, Tocris, UK) and recorded for another 60-120 minutes. Data was averaged over -35 to -5 minutes for baseline measurements and 30 to 60 minutes for on-drug measurements (relative to drug injection). The experiment was repeated after at least 48 hours of rest (see Figure 1A).

**Figure 1.**
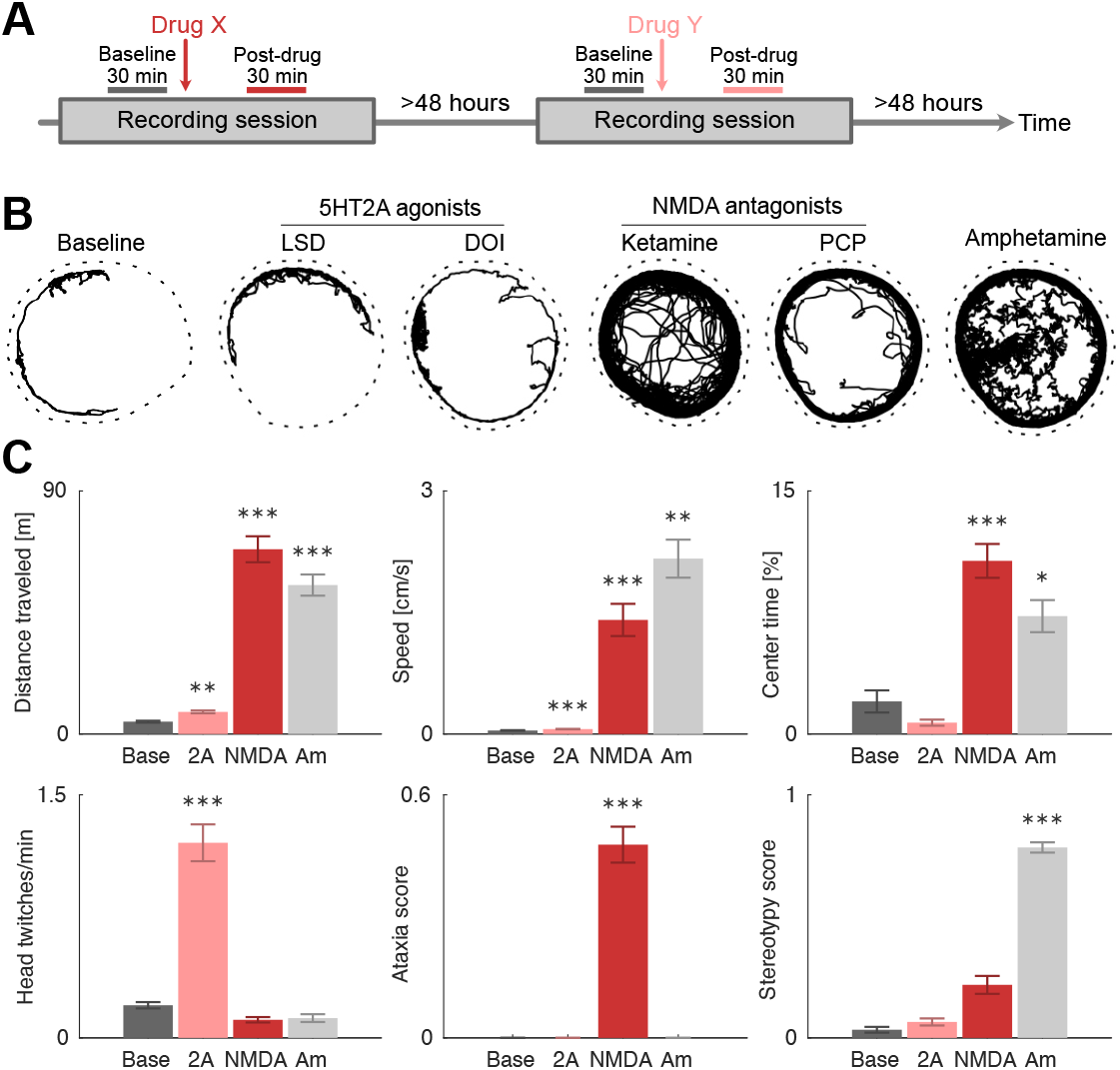
A specific pattern of behavioral changes is induced by each drug class. **A.** Timeline of experiment. Each recording session consisted of 60 min baseline followed by a drug injection and recording for another 60-120 min. Behavioral and electrophysiological data were averaged over -35 to -5 minutes for baseline measurements and 30 to 60 minutes for on-drug measurements (relative to drug injection). At least 48 hours passed between recording sessions. **B.** Examples of tracked motion for each condition. On baseline, the animal was mostly passive and moved occasionally in bouts along the walls of the circular arena (indicated by the dashed line). On the classic psychedelics LSD and DOI, the locomotion behavior was very similar to baseline. In contrast, the dissociative psychedelics ketamine and PCP induced clear hyperlocomotion, and especially ketamine induced ataxic, unstable gait. Amphetamine induced strong hyperlocomotion and vigorous sniffing (seen here as wiggly traces). **C.** Average changes in behavior for each condition (Base = baseline, 2A = LSD or DOI, NMDA = ketamine or PCP, Am = amphetamine). Bars show mean and SEM, asterisks show significance at the p<0.05 (*), p<0.01 (**) and p<0.001 (***) levels (nested ANOVA). Top let: distance traveled during 30 min. Top center: speed. Top right: percentage of time spent in the center. Bottom let: number of head-twitch responses per minute. Bottom center: ataxia score

### Signal acquisition

Local field potential (LFP) and single unity activity were recorded with the Neuralynx multichannel recording system using a unity gain preamplifier (Neuralynx, MT, USA) or with the OpenEphys acquisition system (Siegle et al., 2017) using 4 Intan RHD2132 amplifier boards with on-board AD-converters and accelerometers (Intan technologies, CA, USA). See Supplementary methods for more details. Synchronized video was recorded at 25 fps with a camera placed above the arena.

### Behavioral scoring

Behavior was scored offline from the videos for 1 minute every 10 minutes. Behaviors were scored from 0 - 3 depending on their prevalence (0 = not present, 1 = present for more than 5 seconds, 2 = present for more than 30 seconds, 3 = present continuously). See Table S3 and Supplementary methods for detailed definitions of the scored behaviors.

### Video tracking

The centroid position of the animal was tracked using algorithms adapted from Santana et al. (2014), and locomotion speed and distance traveled was then calculated. A center area was defined as a circle with 2/3 the radius of the arena and the fraction of time spent in the center was quantified.

### Head-twitch response

Head-twitch responses (HTR) were detected using the on-board accelerometers on the Intan RHD2132 amplifier boards together with custom-made Matlab code available at https://github.com/NRC-Lund/htrdetector. See Figure S3 and Supplementary methods for more details. Neuralynx recordings lacked accelerometer data and were analyzed manually from the videos.

### Spike sorting

Extracted spikes were clustered according to a hierarchical clustering scheme using a 2 ms refractory period (Fee et al., 1996). See Supplementary methods for more details. Units were classified into putative cell types based on waveform features as previously described (Halje et al., 2012). A neuron remained unclassified if the probability of belonging to a cell type was <0.75 for both types. Figure S4 and Table S4 show the clustering and summarize the waveform features.

### LFP power spectral densities

To emphasize local sources of the measured electrical potential, bipolar LFP time series were computed from all unique pairs of electrodes from the same structure. Power spectra for truly rhythmic activity was separated from arrhythmic “fractal” activity using the IRASA method (Wen & Liu, 2016). Least-square fitting of each spectrum to the function

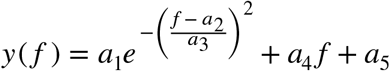

allowed us to define peak height (*a*_1_) and peak frequency (*a*_2_) parametrically and to define criteria for when a peak was detected (see Figure S7 and Supplementary methods for more details).

### Instantaneous phase and amplitude

To quantify the instantaneous phase and amplitude of HFOs, the analytical signal was calculated (Matlab *hilbert* function) from monopolar LFP time series after bandpass filtering ±5 Hz around the median HFO frequency of each recording. Amplitude auto- and cross-correlations were calculated from the instantaneous amplitude time series. Mean phase differences and resultant vector lengths were calculated using the CircStat toolbox (Berens, 2009). A wire pair was said to be phase inverted if Δ*φ* | > 3*π* /4 and if *κ* > 1, where *Δφ* is the average phase difference and κ is the concentration parameter of the von Mises distribution.

### Granger causality

Granger causality was calculated from bipolar LFP time series using the *one_bi_ga*.*m* function of the BSMART toolbox (Cui et al., 2008) with a 500 ms window length and a model order of 5. Spectral peaks were detected using the Matlab *findpeaks*.*m* function with default settings. Only the highest peak was analyzed further if multiple peaks were detected. The total Granger causality between regions was estimated as the median of the amplitude of the Granger causality peaks from all relevant wire pairs.

### Statisical analysis

Nested ANOVA (Aarts et al., 2014) was used unless otherwise stated. Please see Supplementary Methods for more details.

## RESULTS

### Behavior

There are known behavioral cues in rodents that predict psychedelic effects in humans (Hanks & González-Maeso, 2013; Hetzler & Swain Wautlet, 1985). However, those cues are different for classic and dissociative psychedelics; 5-HT2A agonists mainly induce head-twitch responses, while NMDA antagonists mainly induce hyperlocomotion and ataxia. Therefore, the first step in our study was to compare the effects of classic and dissociative psychedelics to the non-psychedelic control on a broad set of behaviors and look for psychedelic-specific behavioral changes.

As expected, the NMDA antagonists induced hyperlocomotion and ataxia, while 5-HT2A agonists induced head-twitch responses and amphetamine induced hyperlocomotion and stereotypy (p<0.001; Figure 1B, 1C, S1 and S2). We found no evidence for a behavior that was similarly and specifically altered by both 5-HT2A agonists and NMDA antagonists. Importantly, this means that psychedelic-specific changes in neuronal activity cannot be explained trivially by changes in motor behavior.

### Unit activity

In total, 365 units were identified from 169 recording sites in 9 freely behaving animals (Figure 2A-C). When comparing the activity of all neurons from all structures to baseline, we observed that the spontaneous firing rates of a large proportion of cells on this global scale were affected by the drugs: 65% were inhibited and only 8% were excited with 5-HT2A agonists, while a more balanced response was found for the NMDA antagonists (40% were inhibited and 40% were excited at the p < 0.01 level; Figures 2D and 2E). As a comparison, in response to amphetamine, 32% were inhibited and 46% were excited (Figure S5).

**Figure 2.**
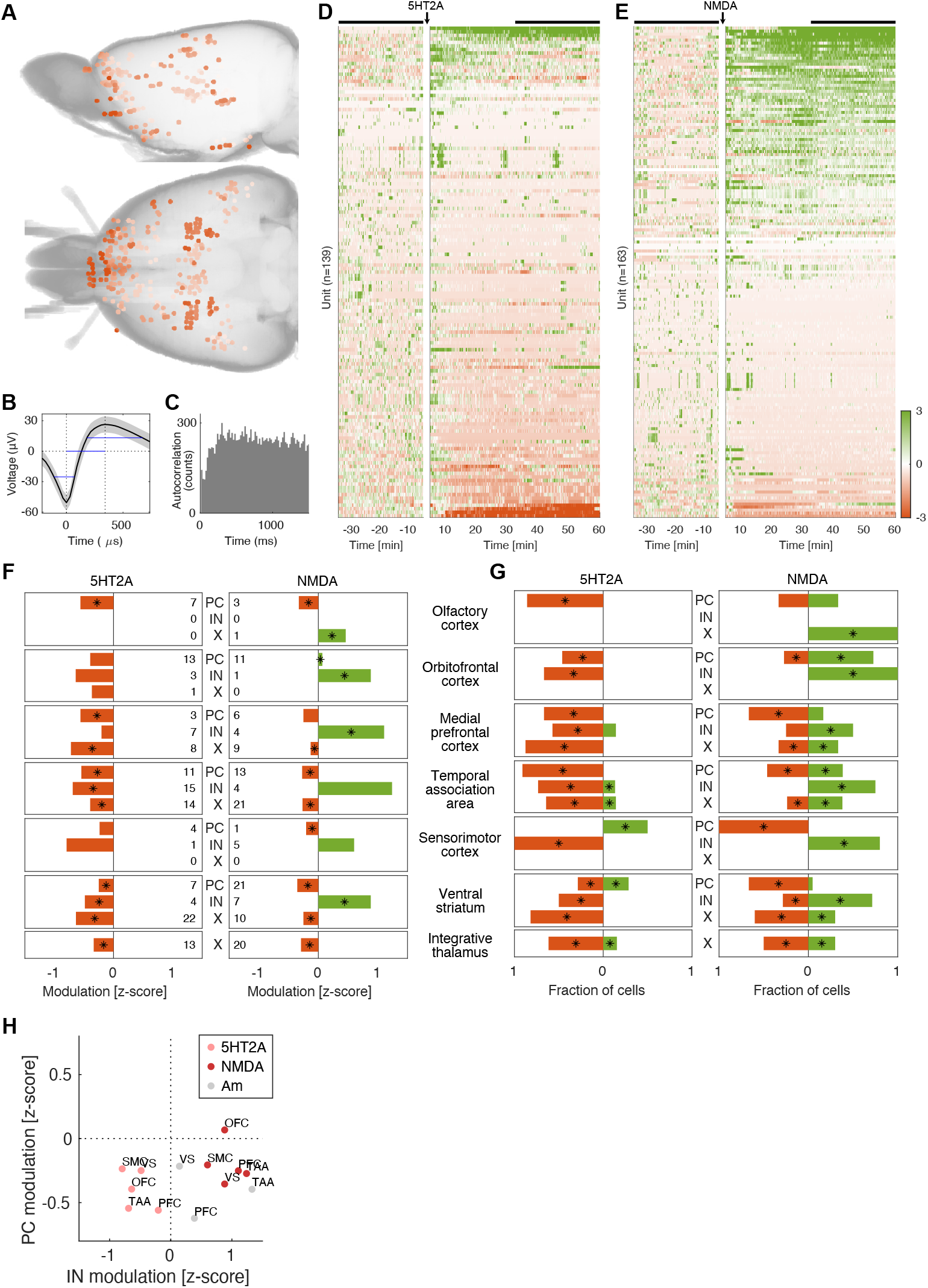
Modulation of neuronal firing rates in response to classic and dissociative psychedelics. **A.** Electrode locations as determined by computer tomography. **B.** Example average waveform from a single unit in prelimbic cortex. The grey area indicates SEM. The blue lines indicate waveform features (peak width, peak-to-valley time and valley width) used to classify the neuron as a putative principal cell. **C.** Spike autocorrelogram of the example unit in B. **D-E**. Standardized neuronal firing rate responses to 5-HT2A agonists (D) and NMDA antagonists (E). Each row shows the activity of a single unit and rows are rank ordered according to the response IN modulation [z-score] during the drug period (indicated by the black bar). **F.** Average standardized neuronal firing rate responses to 5-HT2A agonists (let) and NMDA antagonists (right) for different cell populations (PC=putative principal cells, IN=putative interneurons, X=unclassified cells). The response is calculated as average z-scores during 30 to 60 minutes post drug injection compared to baseline (-35 to -5 minutes). Asterisks indicate significance at the p<0.05 level (nested ANOVA). The numbers next to the cell labels indicate the number of cells in each population. **G.** Fraction of modulated cells as response to 5-HT2A agonists (left) and NMDA antagonists (right) for different cell populations (PC=putative principal cells, IN=putative interneurons, X=unclassified cells). The fractions of downmodulated cells are shown in red and upmodulated cells are shown in green. Asterisks indicate significance at the p<0.05 level (binomial test). **H.** Comparison of IN vs PC rate modulations reveals a similar pattern of modulation for NMDA and amphetamine (IN inhibition, PC excitation), while 5-HT2A induces inhibition in both IN and PC populations. 5-HT2A=pink, NMDA=red, amphetamine=grey. OFC=orbitofrontal cortex, PFC=medial

We further investigated the specific effects of each drug on different cell populations. After grouping of the recording sites based on structural and functional similarity (Table S1), 7 groups had enough cells to warrant further analysis of population modulation (≥4 cells from both 5-HT2A and NMDA experiments; Table S5); olfactory cortex, orbitofrontal cortex, medial prefrontal cortex, temporal association area, sensorimotor cortex, ventral striatum and integrative thalamus. Cell classification based on waveform features was performed on all these structures except thalamus, allowing for a putative division into principal cells (PC), interneurons (IN) and an intermediate group of cells that could not reliably be assigned to either group (unidentified cells; X) (see Figure S4).

For the 5-HT2A agonists the dominating effect on firing rates was inhibition in each of these structures, both in terms of net standardized rates and number of significantly up/down modulated cells (see Figure 2F-G, left panels). For the NMDA antagonists the effect on firing rates was more mixed, with many populations responding with both inhibition and excitation (see Figure 2G, right panel). However, in terms of net standardized rates, we observed that INs were excited and PCs were inhibited in most structures (see Figure 2F, right panel).

To look for rate modulations specific to the psychedelic state, we grouped all types of psychedelic drugs and compared the firing rates to the non-psychedelic control compound amphetamine. Only neurons in integrative thalamus showed a significant difference in this comparison, with a modest reduction of -0.3 (z-scored) in the psychedelic state compared to an increase of 1.6 in the amphetamine state (p < 0.05). Further, the rate modulations for 5-HT2A and NMDA were not significantly different in integrative thalamus (we also controlled that the baselines for all drugs were statistically similar; p > 0.05). However, on a systems level the NMDA modulations throughout all the recorded structures resembled the amphetamine modulations more than the 5-HT2A modulations. This is illustrated by Figure 2H where IN modulation is plotted against PC modulation for each structure.

NMDA and amphetamine had a similar pattern of simultaneous IN excitation and PC inhibition, while 5-HT2A had simultaneous inhibition in both IN and PC. The relative dissimilarity of the clusters was quantified by measuring their Bhattacharyya distance (Kailath, 1967): 5HT2A-NMDA was the most dissimilar pair with a distance of 6.6, while NMDA-amphetamine was the most similar pair with a distance of 0.43. The distance for the 5HT2A-amphetamine pair was 0.98.

In summary, 5-HT2A agonists caused widely distributed inhibition in both principal cells and interneurons. NMDA antagonists caused similar inhibitory effects in principal cells, but opposite, excitatory effects in interneurons. A psychedelic-specific modulation was found in integrative thalamus, but on a systems level the effects of the NMDA antagonists resembled the effects of amphetamine more than those of the 5-HT2A agonists.

### LFP oscillations

To extend on previous results on psychedelics-induced HFOs, we recorded simultaneously from several cortical and subcortical regions and characterized the extent of HFOs, as well as the interactions between HFOs in different regions.

#### Extent and prevalence of HFOs

As expected, both 5-HT2A agonists and NMDA antagonists consistently caused strong increases in HFOs around 150 Hz, as exemplified in Figure 3A and 3B. HFOs increased both in terms of their amplitude (Figure 3C) and their prevalence (as measured by the detection rate; Figure 3D and Figure S7). In contrast, amphetamine increased broad gamma activity (30-80 Hz) without causing HFOs (Fig. 3B, bottom, Figure 3D and Figure S6).

**Figure 3.**
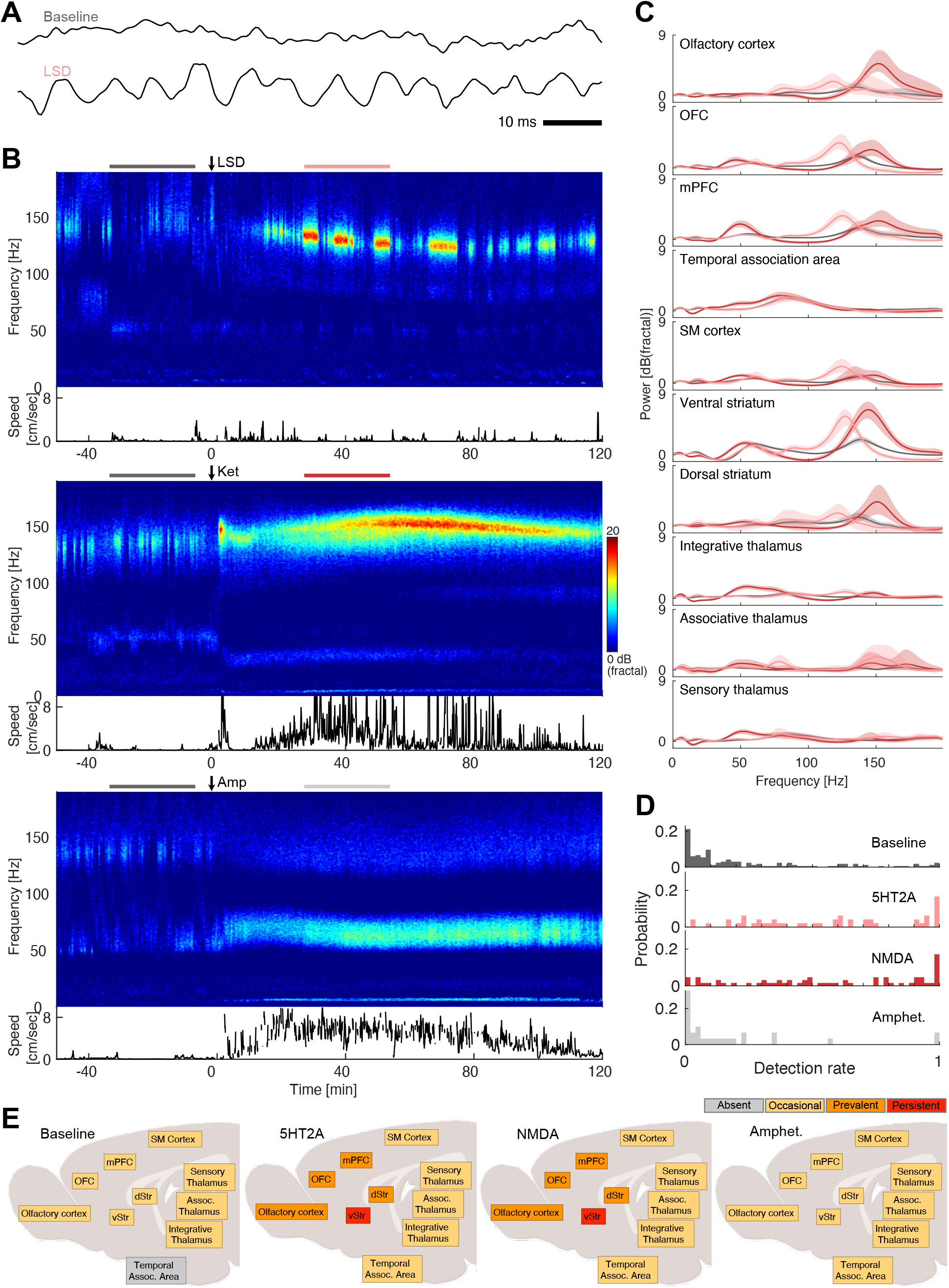
HFOs are enhanced by classic and dissociaive psychedelics but not by amphetamine. **A.** Example traces of LFPs recorded in the ventral striatum (shell of nucleus accumbens) before (top) and after (bottom) LSD administration. The scale bar indicates 10 ms. The signal was low pass filtered at 500 Hz. **B.** LFP spectrograms from the ventral striatum during administration of ketamine (top), LSD (middle) and amphetamine (bottom). A clear increase in high-frequency oscillations around 150 Hz (HFOs) is evident after injection of ketamine or LSD, but not after amphetamine. The color scale is in units of dB_fractal_, i.e. decibels normalized to the fractal noise background. The translational movement speed is shown below each spectrogram. **C.** LFP power spectra averaged over time and treatment groups (baseline=grey, 5-HT2A agonists=pink, NMDA antagonist=red). The time periods used were -35 to -5 minutes for baseline and 30 to 60 minutes for drug treatment relative to injection (also indicated in panel B as horizontal bars above each spectrogram). **D.** Distribution of HFO detection rates in different structures, showing that high detection rates are much more common in 5-HT2A and NMDA conditions. Each value is the average detection rate for a condition in a structure during one recording session. **E.** Summary of HFO detection rates in different structures, showing a similar pattern of HFO prevalence for 5-HT2A agonists and NMDA antagonists. HFOs were classified as persistent (red; more than 90% detections in more than 33% of sessions), prevalent (orange; more than 50% detections in more than 33% of sessions), occasional (yellow, if not in any other class) or absent (grey; more than 5% detections in less than 5% of sessions).

In data pooled from all structures and individuals there was a clear shift towards higher HFO detection rates in 5-HT2A and NMDA conditions, but there was also a large spread in the detection rate distributions (Figure 3D), indicating variation across structures. To further illustrate the extent and prevalence of HFOs in different structures we defined 4 prevalence classes based on the distribution of detection rates (see Figure S8); for each structure and condition, HFOs were classified as persistent (>90% detections in >33% of sessions), prevalent (>50% detections in >33% of sessions), occasional (if not in any other class) or absent (>5% detections in <5% of sessions). According to this classification, 5-HT2A agonists and NMDA antagonists increased HFO prevalence in an identical pattern, while the pattern for the non-psychedelic compound amphetamine was very similar to the baseline pattern (Figure 3E).

#### HFO dynamics and interactions between structures

Next, we addressed the question if psychedelics-induced HFOs are generated in a single structure and propagate to other regions, or whether they are generated in a more distributed fashion. We did this by examining how HFOs in different structures were related in terms of their amplitude, frequency and phase.

Qualitatively, the HFOs had a spindle-like amplitude modulation and strong co-modulation within structures, while co-modulation between structures seemed to be weaker and more complex (Figure 4A). The autocorrelograms of the instantaneous amplitudes were consistent with a typical spindle length of about 50 ms (the autocorrelation function had a smooth peak with FWHM=50±18 ms; Figure 4B). In some electrodes, the autocorrelograms revealed periodic amplitude modulation at the theta frequency (also seen in Figure 4B). Cross-correlograms of the instantaneous amplitudes confirmed that the amplitude modulation was generally more strongly correlated within structures than between structures (median correlation coefficient 0.82±0.10 and 0.40±0.15, respectively; p<0.001, Wilcoxon rank sum; Figure 4C). However, some pairs of structures showed co-modulations that were similar in strength to the within-structure values (see the group above 0.7 in Figure 4C). These strong amplitude correlations were found between olfactory cortex, ventral striatum and orbitofrontal cortex, and between medial prefrontal cortex and ventral striatum.

**Figure 4.**
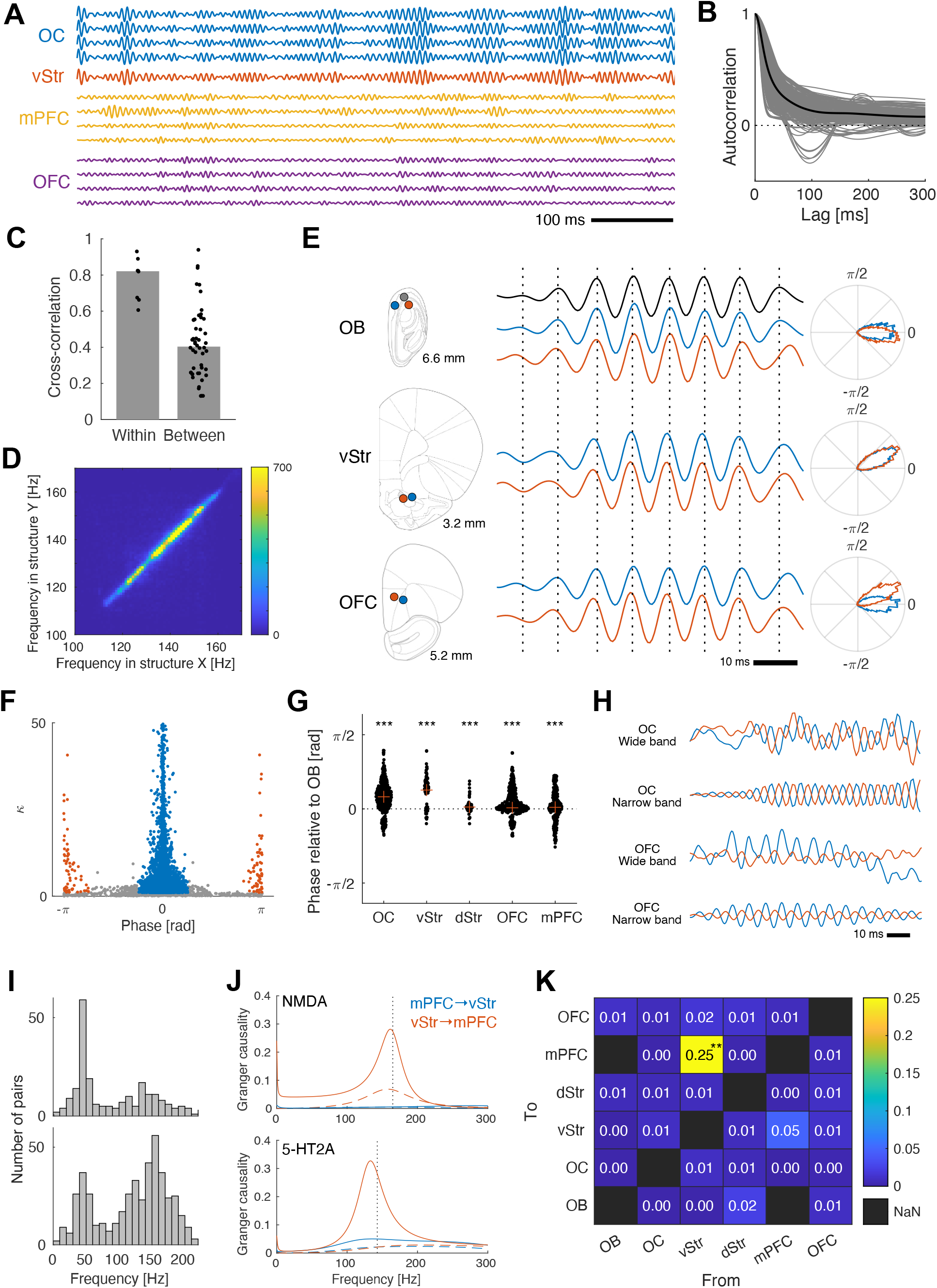
HFOs are globally phase locked but have multiple sources. **A.** Example traces of bandpass filtered LFPs (110-190 Hz, zero-phase FIR) recorded synchronously from the olfactory cortex (OC; blue), the ventral striatum (vStr; red), the medial prefrontal cortex (mPFC; yellow) and the orbitofrontal cortex (OFC; purple) during LSD treatment. Strong co-modulation of the HFO amplitude can be seen both within and between structures, as well as examples of independent modulation within and between structures. **B.** Autocorrelograms of the instantaneous amplitude of all channels with clear HFOs during the psychedelic state (5-HT2A or NMDA). The black line is the average. Most autocorrelograms had a single clear peak (FWHM=50±18 ms) consistent with spindle-like amplitude modulation with a spindle length of about 50 ms. **C.** Cross-correlations of the instantaneous amplitude between all channel pairs with clear HFOs during the psychedelic state (5-HT2A or NMDA). Pairs with both channels in the same anatomical structure (“Within”) had higher cross-correlations on average than pairs with channels in different structures (“Between”). However, some pairs showed between-structure co-modulations that were similar in strength to the within-structure values. **D.** 2D histogram showing the relationship between HFO frequencies in pairs of structures. Each data point comes from an 8 s time window. The high count on the diagonal shows that the frequency is very similar in all structures at any given time, despite a high degree of frequency modulation. **E.** Example traces showing a single HFO spindle recorded synchronously from 7 electrodes in the olfactory bulb (OB; top), the ventral striatum (VS; middle) and the orbitofrontal cortex (OFC; bottom) during treatment with LSD. Electrode positions are shown to the left. Vertical dashed lines are aligned to the peaks of the top OB trace (black) to facilitate comparisons of peak times between electrodes. Polar plots to the right show histograms of the phase difference of each electrode relative to the black OB electrode (based on the whole drug treatment period 30-60 minutes after injection). **F.** Scatter plot showing mean phase differences and κ values for the phase difference distributions of each electrode pair (n=6237). Most pairs had a non-random phase relationship (86% with *κ*>1). Of those, 95% had a mean phase difference close to 0 (*φ* | < *π* /4 ; blue dots) and 2% had an inverted phase (*φ* | > 3*π* /4 ; red dots). **G.** Swarm plot of phase differences relative to the olfactory bulb for all electrode pairs (n=1686) grouped on structure. Each black dot is one electrode pair and red crosses indicate the median for the structure. A positive value means that the structure leads the olfactory bulb. Asterisks indicate that medians are significantly different from zero at the p<0.05 (*), p<0.01 (**) and p<0.001 (***) levels (Wilcoxon signed rank). OB=olfactory bulb, OC=olfactory cortex, vStr=ventral striatum, dStr=dorsal striatum, OFC=orbitofrontal cortex, mPFC=medial prefrontal cortex. **H.** Examples of phase inversion in LFP traces from two nearby electrodes in the anterior olfactory nucleus (AON) and two nearby electrodes in the orbitofrontal cortex (OFC). This indicates the presence of local current dipoles between each electrode pair. Both raw LFP traces (“Wide”) and bandpass filtered traces (“Narrow”) are shown. Note that the AON pair is not recorded synchronously with the OFC pair in this example. **J**. Example Granger causality spectra for one bipolar measurement in medial prefrontal cortex (mPFC) and one bipolar measurement in ventral striatum (vStr) during an NMDA antagonist experiment (top) and a 5-HT2A experiment (bottom). The blue spectra show the causality of mPFC on vStr, while the red spectra show the causality in the opposite direction. Solid lines show the drug treated period and dashed lines show the corresponding baselines. The dotted vertical lines indicate the HFO frequency in the corresponding recording. **K**. Median Granger causality values (calculated from the spectrum peak in the HFO band) for all co-recorded structures with clear HFOs. Black squares indicate missing data. Asterisks indicate that medians are significantly different from zero at the p<0.05 (*), p<0.01 (**) and p<0.001 (***) levels (Wilcoxon signed rank). Only vStr→mPFC showed a causality significantly above zero.

The oscillation frequency was remarkably similar in different brain structures when comparing simultaneous oscillations at different sites, despite large variations between individuals and between conditions (see Figure 4D). Such frequency co-modulation could occur if a single dominant source of oscillating firing rates entrains all other structures via synaptic transmission. This may be considered a likely scenario given previous data showing the importance of the olfactory bulb in the generation of HFOs (Hunt et al., 2019; Wróbel et al., 2020). However, further analyses of phase relationships did not support this hypothesis: Figure 4E shows a single HFO spindle recorded simultaneously from 7 electrodes in the olfactory bulb, the ventral striatum and the orbitofrontal cortex. In this example, the HFOs in the olfactory bulb had almost zero phase difference, while the HFOs in the ventral striatum led the olfactory bulb by about 0.5 radians. Orbitofrontal HFOs had similar phases, ranging between 0 and 0.5 radians in this example. These small – but often non-zero – phase differences were a general finding: In electrode pairs with detectable HFOs (median amplitude >5 μV, n=6237), most pairs had a non-random phase difference (86% with kappa>1), and 95 % of those pairs had an absolute phase difference smaller than pi/4 (see the blue group in Figure 4F). When we compared the phase of the olfactory bulb to different structures, we saw small deviations from zero in all structures (ranging from 0.001 to 0.45 radians, corresponding to temporal delays of <1 ms; see Figure 4G).

The observed near-zero phase lags are not consistent with a single source propagating synaptically or via volume conduction. An alternative explanation is that the HFOs are generated by a system of several self-sustaining but weakly interacting oscillators located in multiple structures. Such systems are known to have stable states with near-zero phase lags; similar to a standing wave (Acebrón et al., 2005). To find support for the presence of local HFO generators outside the olfactory bulb, we looked for phase inversions in measurements from adjacent electrodes, since an inverted phase indicates that a local current dipole is present between the electrodes. Such inversions were indeed found in about 2% of intrastructural electrode pairs (defined as an absolute phase difference larger than 3π/4 and *κ* ; see the red group in Figure 4F) and in several structures, including the olfactory bulb, the olfactory cortex, the orbitofrontal cortex, the medial prefrontal cortex and the ventral striatum (Figure 4H).

To further map the network of influences between structures, we calculated the Granger causality between structure pairs in the frequency domain. The Granger causality spectra often had clear peaks either in the HFO band or in the gamma band. During baseline, the peak of the Granger causality was mostly in the low gamma band, while it was mostly in the HFO band during the psychedelic state (Figure 4I-J). The mean Granger causality in the HFO band (HFO frequency ±10 Hz) increased by 97% compared to baseline (p<0.001, Wilcoxon rank sum), while it was not significantly changed in the gamma band (25-75 Hz; p=0.34). A structure-by-structure analysis revealed that the Granger causality was clearly strongest from ventral striatum to mPFC (Figure 4K).

In summary, the most specific neurophysiological correlate of psychedelic drug action was the enhancement of widespread phase-synchronized HFOs around 150 Hz. In the psychedelic state, we have identified a network consisting of the olfactory bulb, the olfactory cortex, the ventral striatum, the orbitofrontal cortex and the medial prefrontal cortex that are tightly coupled in the HFO band.

## DISCUSSION

Several models propose explanations for how the known pharmacological effects on single neurons result in changes at the neuronal systems level, and how these in turn are related to subjective psychedelic experiences (Carhart-Harris & Friston, 2019; Doss et al., 2022; van Elk & Yaden, 2022; Vollenweider & Preller, 2020). Some models explicitly state a direct link between firing rates and functional changes, like the thalamocortical gating model, in which increased mPFC excitability disinhibits the thalamus and reduces its ability to gate sensory information (Vollenweider & Geyer, 2001). Other models have indirectly linked the activation of certain cell populations to the disintegration of canonical network states, as seen for example in the reduced BOLD correlation between nodes in the default mode network (Carhart-Harris et al., 2016). Existing models have however highlighted apparently contradictory evidence and the field has suffered from an almost complefte lack of *in vivo* electrophysiology data from awake animals that could link the pharmacological effects on single cells with changes in information processing in the brain as a whole (Smausz et al., 2022).

With simultaneous large-scale microwire recordings in multiple structures in freely behaving animals, we here show that different classes of psychedelics affect firing rates differently, while they cause similar changes in population dynamics in the form of aberrantly strong HFOs. We also show that the HFOs facilitate functional coupling between brain structures, both in terms of phase synchronization and Granger causality. This could have major effects on the exchange and integration of information across these neuronal systems.

Our results cast doubts on models suggesting that specific changes in firing rates are directly linked to the psychedelic state, since the appearance of HFOs is largely independent of population firing rates. This is a surprising finding given the dominant hypothesis that both classic and dissociative psychedelics exert their effect via increased release of glutamate in cortico-limbic circuits (de Gregorio et al., 2018). Intuitively, increased glutamate release should lead to increased excitation. However, biological neuronal networks have several homeostatic mechanisms to regulate the balance between excitation and inhibition, which makes it difficult to predict how changes in glutamate signaling affect the overall behavior of the network. Indeed, the seminal studies reporting increased AMPA-dependent excitatory currents in pyramidal cells did not observe simultaneous increases in firing rates of those neurons (Aghajanian & Marek, 1997, 1999; Zhou & Hablitz, 1999) and, more generally, it is well known that psychedelics are not proconvulsant.

Perhaps it is fruitful to focus less on the excitatory role of glutamate, and more on how it may change the strength and temporal dynamics of effective synaptic coupling. Theoretical work has shown that the strength and dynamics of synaptic coupling play crucial roles in determining network behavior and, in particular, if a network has stable periodic states (Richter et al., 2013). Intriguingly, both classic and dissociative psychedelics decrease NMDA-mediated synaptic currents (Arvanov et al., 1999), while they increase AMPA-mediated currents (as mentioned above). However, AMPA channels have a much shorter deactivation time constant than NMDA channels (AMPA: 2-5 ms, NMDA: 50-100 ms), which will lead to dramatically faster temporal dynamics of the excitatory postsynaptic currents. This decreased “temporal smoothing” could in turn enable stable oscillatory states with a shorter period than would otherwise be possible.

While this might be a possible explanation for the appearance of local oscillatory states, the long-range HFO synchronization observed in the current study is still perplexing. Intuitively, it seems unlikely that such fast oscillations can synchronize across long distances considering the sizeable delays caused by the propagation of action poftentials and the delayed activation of chemical synapses. On the other hand, gap junctions and ephaptic coupling could influence neighboring neurons almost instantaneously, but have very short range. However, mathematical analysis of idealized coupled oscillators has shown that stable synchronous states can exist with only local connectivity and even with delayed influences (Acebrón et al., 2005; Gerstner, 1996). Interestingly, such systems often display a surprising complexity, where multiple stable synchronous states can co-exist and have different synchronization frequencies (Yeung & Strogatz, 1999).

Taken together, the current study represents a first step towards bridging psychedelics-induced physiological phenomena at the single cell level, via networks, to global brain states. Increasing our mechanistic understanding of how this class of substances induce psychedelic states will be essential to develop improved therapies for several neuropsychiatric conditions.

## ACKNOWLEDGEMENTS

Lund University Bioimaging Centre (LBIC), Lund University, is gratefully acknowledged for providing experimental resources. The IRASA computations were enabled by resources provided by the Swedish National Infrastructure for Computing (SNIC) at LUNARC.

## AUTHOR CONTRIBUTIONS

I.B. designed and performed experiments, and contributed to manuscript writing;

S.B. performed experiments and contributed to manuscript writing.

E.W. performed behavioral analysis and assisted during experiments.

J.W. contributed to spike analysis and assisted during experiments.

P.P. contributed to manuscript writing and experimental design.

P.H. designed experiments, performed analysis and wrote the paper.

## FUNDING

The study was supported by grants from BABEL (Erasmus Mundus), Crafoord Foundation, Insamlingsstitelserna, Kempe Foundation, Barncancerfonden, Kocks Foundation, Kungliga Fysiografiska Sällskapet, Magnus Bergvall Foundation, Olle Engkvist Foundation, Oskarfonden, Parkinsonfonden, Petrus och Augusta Hedlunds Stifelse, Promobilia, Segerfalk Foundation, Sigur & Elsa Goljes Minne Foundation, Sven-Olof Jansons livsverk, Svenska Sällskapet för Medicinsk Forskning (SSMF), Thurings Foundation, Umeå Universitet, Vetenskapsrådet (#2018-02717 and #2016-07213) Wenner-Gren Foundation and Åhlén Foundations.

## COMPETING INTERESTS

The authors declare that they have no competing interests.

## Notes

### Competing Interest Statement

The authors have declared no competing interest.

