## Supplementary Methods, Tables and Figures for "Classic and dissociative psychedelics induce similar hyper-synchronous states in the cognitive-limbic cortex-basal ganglia system"

for

This PDF includes:

- Supplementary Methods
- Supplementary Tables S1-S5
- Supplementary Figures S1-S8

### Animals

Nine Sprague-Dawley rats were used (Taconic, Denmark) in this study. Animals were kept in plastic cages on a 12:12-h light-dark cycle (lights on from 5:30 h to 17:30 h). All procedures were approved in advance by the Malmö/Lund ethical committee of animal experiments.

### Construction of electrode arrays

Microelectrode arrays were built and implanted as previously described (Ivica et al., 2014), and allowed us to record local field potentials (LFP) and single units from up to 128 channels, for several weeks in freely moving animals. Briefly, formvar-insulated tungsten wires (33 and 50 µm diameter, California Fine Wire Co., CA) were arranged and cut to target up to 7 regions bilaterally: olfactory, orbitofrontal, sensorimotor, and prefrontal cortices, the hippocampus, ventral striatum and mediodorsal thalamus. The wires were soldered to a connector on a custom-made circuit board and the whole ensemble was fixated with UV curing.

### Implantation surgery

Implantation surgeries were performed under fentanyl/medetomidine anesthesia (0.3/0.3 mg/kg, i.p.) and followed the procedures previously described (Ivica et al., 2014). Using a micromanipulator, the electrodes arrays were implanted in both hemispheres, and fixated with dental acrylic attached to 5 - 8 screws in the skull. A 200 µm thick silver wire was attached to three of these screws located on the occipital bone and was used as a ground connection from the animal to the recording system. After the implantation, the anesthesia was reversed by atipamezole hydrochloride (5 mg/kg, i.p.). Saline and the postoperative analgesic buprenorphine (0.5 mg/kg, subcutaneous injection) were administered. The animals were allowed to recover for at least two weeks, during which they were daily monitored and had their diet supplemented with a nutrient fortified water gel when needed (DietGel® Recovery, Karlslunde, Denmark).

### Verification of electrode positions

Electrode positions were verified post mortem in 5 animals as follows: Animals were anesthetized with a lethal dose of sodium pentobarbital (100 mg/kg) and transcardially perfused with saline 0.9% (room temperature) followed by paraformaldehyde 4% (4 °C). Heads were stored in paraformaldehyde at 4 °C until scanned with computed tomography (CT) on a MILabs XUHR system (MILabs, Netherlands; 65 kV peak energy, 0.13 mA current, 90 ms exposure time, 200 microns Cu filter, 100 microns Al filter). The head was oriented so that the electrode wires were perpendicular to the photon beam to minimize artifacts. CT volumes were reconstructed with a voxel size of 20 microns and wire tips were identified semi-automatically in the CT volumes using custom-made software (https://github.com/NRC-Lund/ct-tools). The volumes were registered to an anatomical atlas (Paxinos & Watson, 2007) using lambda and bregma as landmarks, but a manual calibration was usually necessary to optimize the alignment between atlas and scan visually. The resulting affine transformation was used to calculate the atlas coordinates of the wire tips from the voxel coordinates (see Figure 2A). Finally, the wire tips were assigned appropriate anatomical labels based on their location in the atlas.

### Pharmacological treatments

To record the behavioral and electrophysiological effects of pharmacological treatments, animals were placed in a round open field arena and the implant was connected to the amplifier boards. After ~60 min of baseline recording, the animal was intraperitoneally injected with LSD (lysergic acid diethylamid, 0.3 mg/kg, Lipomed AG, Switzerland), DOI (2,5-dimethoxy-4-iodoamphetamine hydrochloride, 2 mg/kg, Lipomed AG, Switzerland), ketamine (Ketaminol, 25 - 50 mg/kg, Intervet AB, Sweden), PCP (phencyclidine hydrochloride, 5 mg/kg, Lipomed AG, Switzerland) or amphetamine (d-amphetamine sulfate, 4 mg/kg, Tocris, UK) and recorded for another 60-120 minutes. Data was averaged over -35 to -5 minutes for baseline measurements and 30 to 60 minutes for on-drug measurements (relative to drug injection). The experiment was repeated after at least 48 hours of rest (see Figure 1A).

### Signal acquisition

Local field potential (LFP) and single unity activity were recorded with the Neuralynx multichannel recording system using a unity gain preamplifier (Neuralynx, MT, USA) or with the OpenEphys acquisition system (Siegle et al., 2017) using 4 Intan RHD2132 amplifier boards with on-board AD-converters and accelerometers (Intan technologies, CA, USA).

For Neuralynx recordings, LFP signals were filtered between 0.1 and 300 Hz, and were digitized at 1017 Hz. Unit activities were filtered between 600 and 9,000 Hz and digitized at 32 kHz. Thresholds for storage of spiking events in each channel were set to 2.5 SDs of the unfiltered signal.

For OpenEphys recordings, wideband signals were digitized and recorded at 30 kHz after bandpass filtering between 0.1 Hz and 10 kHz. LFPs were extracted offline by low pass filtering ($8^{th}$ order Butterworth at 500 Hz) and downsampling to 2000 Hz. Spike waveforms were extracted offline by thresholding at 4 SDs after bandpass filtering between 600 Hz and 9000 Hz ($128^{th}$ order FIR) and extracting 1 ms before and 2 ms after the threshold crossing event. The detector had a dead period of 1 ms. In addition, 3-axis accelerometer data was digitized and recorded at 30 kHz from all 4 amplifier boards.

Video was recorded at 25 fps with a camera placed above the open field arena. It was synchronized to the electrophysiology system using a Master-8 pulse generator (AMPI, Israel).

### Behavioral scoring

Behavior was scored offline from the videos for 1 minute every 10 minutes. Behaviors were scored from 0 - 3 depending on their prevalence (0 = not present, 1 = present for more than 5 seconds, 2 = present for more than 30 seconds, 3 = present continuously). The following behaviors were scored: being still, grooming, rearing, sniffing upwards, sniffing downwards, head-swaying, moving backwards, intermittent turning, unstableness, falling over, lying down and crawling. See Table S3 for more detailed definitions of the scored behaviors. Scores of head-sway, unstableness, falling over, lying down and crawling were averaged to an ataxia score. Similarly, scores of sniffing upwards, sniffing downwards and moving backwards were averaged to a stereotypy score (David Sturgeon et al., 1979).

### Video tracking

Object tracking was performed using algorithms in Matlab adapted from (Santana et al., 2014). Briefly, the foreground was separated from the static background using luminosity thresholding and foreground blobs were tracked between frames using a Kalman filter. The blob belonging to the animal (as opposed to the cable, for example) was identified based on shape parameters. The position of the animal was defined as the blob centroid.

Locomotion speed was calculated based on the translation of the blob centroid during a 1 second window. The distance traveled was calculated as the sum of the speed time series. Normally, rats prefer to stay along the arena walls and the time spent in the center was quantified and interpreted as a measure of disorientation or an increased drive to explore. The center area was defined as a circle with 2/3 the radius of the arena.

### Head-twitch response

Head-twitch responses (HTR) were detected using the on-board accelerometers on the Intan RHD2132 amplifier boards that were attached to the dorsal side of the head via rigid adaptors. The mediolateral acceleration signal was downsampled to 200 Hz and bandpass filtered forwards and backwards with a finite impulse response filter (passband 8-32 Hz, filter order 100). A HTR index was constructed by convoluting the absolute value of the filtered signal with a Gaussian window ($\sigma$=50 ms). HTR events were extracted by detecting peaks in the HTR index that were higher than the threshold (0.4 g) and were separated by more than the detector dead time (1 sec; see Figure S3).

The method was validated with manual inspection of the videos in 23 recordings. The validation resulted in a true positive rate of 95% with a false positive rate of 0.1 %. The method was implemented in the Matlab programming language and is available at https://github.com/NRC-Lund/htrdetector.

Neuralynx recordings lacked accelerometer data and were analyzed manually from the videos.

### Spike sorting

Extracted spikes were dejittered and clustered according to a hierarchical clustering scheme using a 2 ms refractory period (Fee et al., 1996). Noise clusters were detected based on the normalized spike density during the refractory period and were removed if >1. Finally, all clusters were manually reviewed to determine if spike waveforms, firing rates, autocorrelation functions and interspike-interval distributions were physiologically plausible. About 12% of the clusters survived the manual review.

Units were classified into putative cell types based on the following waveform features: valley width, peak width, and peak-to-valley time. The widths were defined as the full width at half maximum (FWHM). The classification was performed by fuzzy k-means clustering with probability of membership >0.75, i.e., units with a probability of membership < 0.75 were labeled as unclassified. Figure S4 and Table S4 shows the clustering and summarizes the statistics of the waveform features for the cell types in all structures.

#### LFP power spectral densities

To emphasize local sources of the measured electrical potential, bipolar LFP time series were computed from all unique pairs of electrodes from the same structure. For each of these time series, spectrograms were calculated with 50%-overlapping 8-s windows (0.12 Hz resolution) using the Irregularly Resampled AutoSpectral Analysis method (IRASA; Wen & Liu, 2016). IRASA separates the arrhythmic, so-called fractal component $S_{\text{fractal}}(f)$ from the power spectrum $S(f)$, and by normalizing the spectrum to the fractal component, it is possible to construct a power spectrum measure that emphasizes truly rhythmic activity:

$$S_{\text{dB(fractal)}} = 10 \log \frac{S(f)}{S_{\text{fractal}}(f)}$$

The spectra $S_{\text{dB(fractal)}}$ from individual electrode pairs were averaged structure-by-structure and fed into a peak detection algorithm that used nonlinear least-square fitting (Matlab `fit` function) to fit each spectrum to the following model:

$$y(f) = a_1 e^{-\left(\frac{f-a_2}{a_3}\right)^2} + a_4 f + a_5$$

This allowed us to quantify peak height ($a_1$) and peak frequency ($a_2$) parametrically, but we also used this model as an oscillation detector by defining a threshold for the goodness-of-fit ($R^2$) and limits for the fitted parameters. Typical conditions for a positive HFO detection were $R^2$>0.2, 2<$a_1$<100 dB, 90<$a_2$<170 Hz, 1<$a_3$<20 Hz, -1<$a_4$<1 and -10<$a_5$<10 (see Figure S7).

#### Instantaneous phase and amplitude

To quantify the instantaneous phase and amplitude of HFOs, monopolar LFP time series were bandpass filtered ±5 Hz around the median HFO frequency of each recording (as determined by the $a_2$ parameter above). We used a 64-order FIR filter (Matlab `fir1` function) backwards and forwards to ensure zero phase lag (Matlab `filtfilt` function). The bandpassed signal was Hilbert transformed into the complex-valued analytical signal $z(t)$ (Matlab `hilbert` function) and instantaneous phase and amplitude could then be calculated as $\varphi(t) = \arg z(t)$ and $r(t) = |z(t)|$.

Amplitude auto- and cross-correlations were calculated from the instantaneous amplitude time series in windows of 60 seconds and were then averaged across windows. For auto-correlations, only channels with a median amplitude above 15 µV were considered. To compare cross-correlations within and between structures, the height of the peak of the cross-correlogram was calculated for each electrode pair.

The mean phase difference between a pair of wires $i$ and $j$ was calculated as $\Delta\varphi_{ij} = \langle \varphi_i(t) - \varphi_j(t) \rangle$, where $\langle\rangle$ denotes the circular mean (function `circ_mean`, Matlab CircStat toolbox (Berens, 2022). Similarly, the resultant vector length $r_{ij}$ was obtained using the function `circ_r` of the same toolbox. The mean phase difference between two brain regions was estimated by averaging all $\Delta\varphi_{ij}$. However, pairs with $r_{ij} < 0.5$ were excluded from the average to ensure that only sufficiently stable phase difference estimates were used. Two wires were said to be phase inverted with respect to each other if $|\Delta\varphi_{ij}| > \frac{3}{4}\pi$ and if $\kappa > 1$, where $\kappa$ is the concentration parameter of the von Mises distribution.

#### Granger causality

Granger causality was calculated from bipolar LFP time series using the `one_bi_ga` function of the BSMART toolbox (Cui et al., 2008) with a 500 ms window length and a model order of 5. Typically, the Granger causality spectrum showed a clear peak in the gamma band or the HFO band. The frequency and amplitude of the peak were detected using the Matlab `findpeaks` function with default settings. Only the highest peak was analyzed further if multiple peaks were detected. The total Granger causality from one region to another region was estimated as the median of the amplitude of the Granger causality peaks from all relevant wire pairs.

### Statistical analysis

Comparisons of behavioral measures were done with a nested ANOVA model using the Matlab `anovan` function (Aarts et al., 2014) with factors State (baseline versus drug), Session and Animal. The factor Session was nested in Animal. Session and Animal were defined as random variables. A separate ANOVA was performed for each drug and for each evaluated parameter. The significance level was set to $\alpha$=0.05.

Comparisons of firing rates were done with a Wilcoxon rank sum test for equal medians (Matlab `ranksum` function) when determining if a unit was significantly modulated compared to its own baseline ($\alpha$=0.01). We used a binomial test (Matlab `binocdf` function) to determine if the number of significantly modulated cells in a population was higher or lower than chance ($\alpha$=0.05). Standardized population rates were compared to baseline with a nested ANOVA model with factors State (baseline vs drug), Session, Animal, Time and Neuron. The factor Session was nested in Animal, while Time was nested in State and Neuron was nested in Session and Animal. Session, Animal and Neuron were defined as random variables and Time was defined as a continuous variable.

HFO amplitude and frequency were compared to baseline with a nested ANOVA model with factors State (baseline vs drug), Session, Animal, Hemisphere and Structure. The factor Session was nested in Animal. Session and Animal were defined as random variables. The model was identical for comparisons between drugs, except that State now had the levels “5HT2A” and “NMDA”, and that Session was nested in both Animal and State. Differences in cross-correlations and Granger causality were tested with the Wilcoxon rank sum test.

### Table S1: Electrodes grouped on anatomical location

Table listing all electrodes grouped on their anatomical location according to the atlas (Paxinos & Watson, 2007). The anatomical locations were further grouped into 10 functional groups. The location of the wires in the “Wires (CT)” column were validated using computer tomography, while the location of wires in the “Wires (no CT)” were inferred from stereotaxic coordinates.

| Abbriviation | Anatomical name | Functional group | Wires (CT) | Wires (no CT) |
|---|---|---|---|---|
| AOE | anterior olfactory nucleus, external part | Olfactory cortex | 1 | |
| AOL | anterior olfactory nucleus, lateral part | Olfactory cortex | 4 | |
| AOP | anterior olfactory nucleus, posterior part | Olfactory cortex | 13 | |
| DEn | dorsal endopiriform nucleus | Olfactory cortex | 8 | |
| IEn | Intermediate endopiriform nucleus | Olfactory cortex | 1 | |
| EPI | external plexiform layer of the olfactory bulb | Olfactory cortex | 2 | |
| Gl | glomerular layer of the olfactory bulb | Olfactory cortex | 5 | |
| GrO | granular cell layer of the olfactory bulb | Olfactory cortex | 2 | |
| Mi | mitral cell layer of the olfactory bulb | Olfactory cortex | 2 | |
| PLCo | posterolateral cortical amygdaloid nucleus | Olfactory cortex? | 1 | |
| Pir | piriform cortex | Olfactory cortex | 20 | |
| AI | agranular insular cortex, ventral part (bordering LO) | Orbitofrontal cortex | 3 | |
| DLO | dorsolateral orbital cortex | Orbitofrontal cortex | 4 | |
| LO | lateral orbital cortex | Orbitofrontal cortex | 31 | 18 |
| MO | medial orbital cortex | Orbitofrontal cortex | 15 | |
| VO | ventral orbital cortex | Orbitofrontal cortex | 15 | |
| FrA | frontal association cortex | Prefrontal cortex? | 8 | |
| IL | infralimbic cortex | Prefrontal cortex | 1 | 12 |
| PrL | prelimbic cortex | Prefrontal cortex | 36 | 62 |
| Au2 | secondary auditory cortex | Temporal association area | 4 | |
| Ect | ectorhinal cortex | Temporal association area | 5 | |
| DLEnt | dorsolateral entorhinal cortex | Temporal association area | 1 | |
| HippD | dorsal hippocampus | Temporal association area | | 16 |
| HippV | ventral hippocampus | Temporal association area | | 94 |
| MoDG | molecular layer of the dentate gyrus | Temporal association area | 8 | |
| Or | oriens layer of the hippocampus | Temporal association area | 9 | |
| opt | olivary pretectal nucleus (bordering hippocampus) | Temporal association area | 1 | |
| M1 | primary motor cortex | Sensorimotor cortex | 11 | |
| M2 | secondary motor cortex | Sensorimotor cortex | 20 | 12 |
| S1 | primary somatosensory cortex | Sensorimotor cortex | 58 | |
| S2 | secondary somatosensory cortex | Sensorimotor cortex | 1 | |
| AcbC | accumbens nucleus, core | Ventral striatum | 1 | 42 |
| AcbSh | accumbens nucleus, shell | Ventral striatum | 8 | 30 |
| CPuV | ventral striatum | Ventral striatum | 1 | |
| Tu | olfactory tubercle | Ventral striatum | 10 | |
| CPuDL | dorsolateral striatum | Sensorimotor striatum | 15 | 8 |
| CPuL | lateral striatum | Sensorimotor striatum | 2 | |
| CM | central medial thalamic nucleus | Integrative thalamus | 1 | |
| LDDM | laterodorsal thalamic nucleus, dorsomedial part | Integrative thalamus | 1 | |
| MD | mediodorsal thalamic nucleus | Integrative thalamus | 6 | 79 |
| PT | paratenial thalamic nucleus | Integrative thalamus | 2 | |
| AD | anterodorsal thalamic nucleus | Associative thalamus | 2 | |
| AM | anteromedial thalamic nucleus | Associative thalamus | 15 | |
| AV | anteroventral thalamic nucleus | Associative thalamus | 13 | |
| MG | medial geniculate nucleus | Sensory thalamus | | 42 |
| Po | posterior thalamic nuclear group | Sensory thalamus | 1 | |
| VG | ventral geniculate nucleus | Sensory thalamus | 2 | |
| VPL | ventral posterolateral thalamic nucleus | Sensory thalamus | 9 | |
| VPM | ventral posteromedial thalamic nucleus | Sensory thalamus | 2 | |
| Rt | reticular thalamic nucleus | Sensory thalamus | 1 | |

#### Table S2: Electrodes grouped on functional group and animal

Table showing the number of wires in each functional group split on animals. Animals ER, ES, FA, FB and FC were CT scanned.

**Count of Location vs. Animal**

| Location \ Animal | EA | EJ | EM | ER | ES | FA | FB | FC | FD |
|---|---|---|---|---|---|---|---|---|---|
| Olfactory cortex | 0 | 0 | 0 | 8 | 2 | 0 | 23 | 26 | 0 |
| Orbital cortex | 0 | 0 | 0 | 6 | 10 | 12 | 21 | 19 | 18 |
| Prefrontal cortex | 19 | 17 | 14 | 8 | 0 | 25 | 12 | 0 | 24 |
| Temporal association area | 23 | 39 | 20 | 0 | 15 | 12 | 1 | 0 | 28 |
| Sensorimotor cortex | 0 | 0 | 0 | 0 | 1 | 33 | 31 | 25 | 12 |
| Ventral striatum | 22 | 21 | 17 | 5 | 9 | 0 | 3 | 3 | 12 |
| Sensorimotor striatum | 0 | 0 | 0 | 0 | 1 | 3 | 11 | 2 | 8 |
| Cognitive thalamus | 23 | 24 | 16 | 0 | 2 | 1 | 6 | 1 | 16 |
| Associative thalamus | 0 | 0 | 0 | 14 | 7 | 1 | 6 | 2 | 0 |
| Sensory thalamus | 10 | 16 | 16 | 8 | 6 | 0 | 1 | 0 | 0 |

#### Table S3: Behaviors assessed by scoring

| Behavior | Definition |
|---|---|
| Being still | Being completely still with all paws touching the ground (cf "Lying down"). This presumably includes both wakeful resting and sleeping. |
| Grooming | Stereotyped rat self-grooming, including paw strokes on the head and body licking. |
| Rearing | Standing up on the hindlimbs. |
| Sniffing upwards | Sniffing and whisking with the head turned upwards. |
| Sniffing downwards | Sniffing and whisking with the head directed forward or toward the ground. |
| Head-swaying | Head swaying from side to side. |
| Moving backwards | Backward locomotion. |
| Intermittent turning | Short locomotion bouts that alternate left and right without resulting in any significant movement forwards. |
| Unstableness | Unstable posture or stumbling, wobbly gait. |
| Falling over | Severe unstableness resulting in falls while moving or standing still. |
| Lying down | Lying down on the side, unable to maintain a normal body posture. Often accompanied by limb movements. |
| Crawling | Movement of limbs with or without change of location while being unable to stand up |

#### Table S4: Summary of spike shape features (mean±SEM μs)

| Structure | Cell type | Peak width | Valley width | Peak-to-valley time |
|---|---|---|---|---|
| Olfactory cortex | PC | 201 ± 25 | 368 ± 13 | 417 ± 11 |
| | IN | 122 ± 31 | 224 ± 31 | 215 ± 5 |
| | X | 175 ± 0 | 276 ± 0 | 325 ± 0 |
| Neocortex | PC | 243 ± 8 | 402 ± 8 | 506 ± 9 |
| | IN | 109 ± 3 | 252 ± 12 | 210 ± 9 |
| | X | 141 ± 6 | 406 ± 12 | 337 ± 7 |
| Temporal association area | PC | 194 ± 13 | 362 ± 11 | 433 ± 11 |
| | IN | 111 ± 4 | 211 ± 12 | 209 ± 10 |
| | X | 166 ± 12 | 276 ± 26 | 333 ± 9 |
| Striatum | PC | 266 ± 7 | 332 ± 7 | 427 ± 7 |
| | IN | 127 ± 6 | 234 ± 9 | 223 ± 15 |
| | X | 185 ± 15 | 254 ± 21 | 361 ± 13 |
| Thalamus | X | 128 ± 6 | 258 ± 17 | 289 ± 20 |

#### Table S5: Number of cells grouped on functional group and drug

Number of cells

| Location | 5HT2A | NMDA | amphet |
|---|---|---|---|
| Associative thalamus | 3 | 2 | 0 |
| Cognitive thalamus | 13 | 20 | 6 |
| Olfactory cortex | 7 | 4 | 0 |
| Orbital cortex | 17 | 12 | 2 |
| Prefrontal cortex | 18 | 19 | 18 |
| Sensorimotor cortex | 5 | 6 | 1 |
| Sensorimotor striatum | 2 | 2 | 0 |
| Sensory thalamus | 1 | 22 | 2 |
| Temporal association area | 40 | 38 | 9 |
| Ventral striatum | 33 | 38 | 25 |

Drug

### Figure S1: Manually scored behaviors

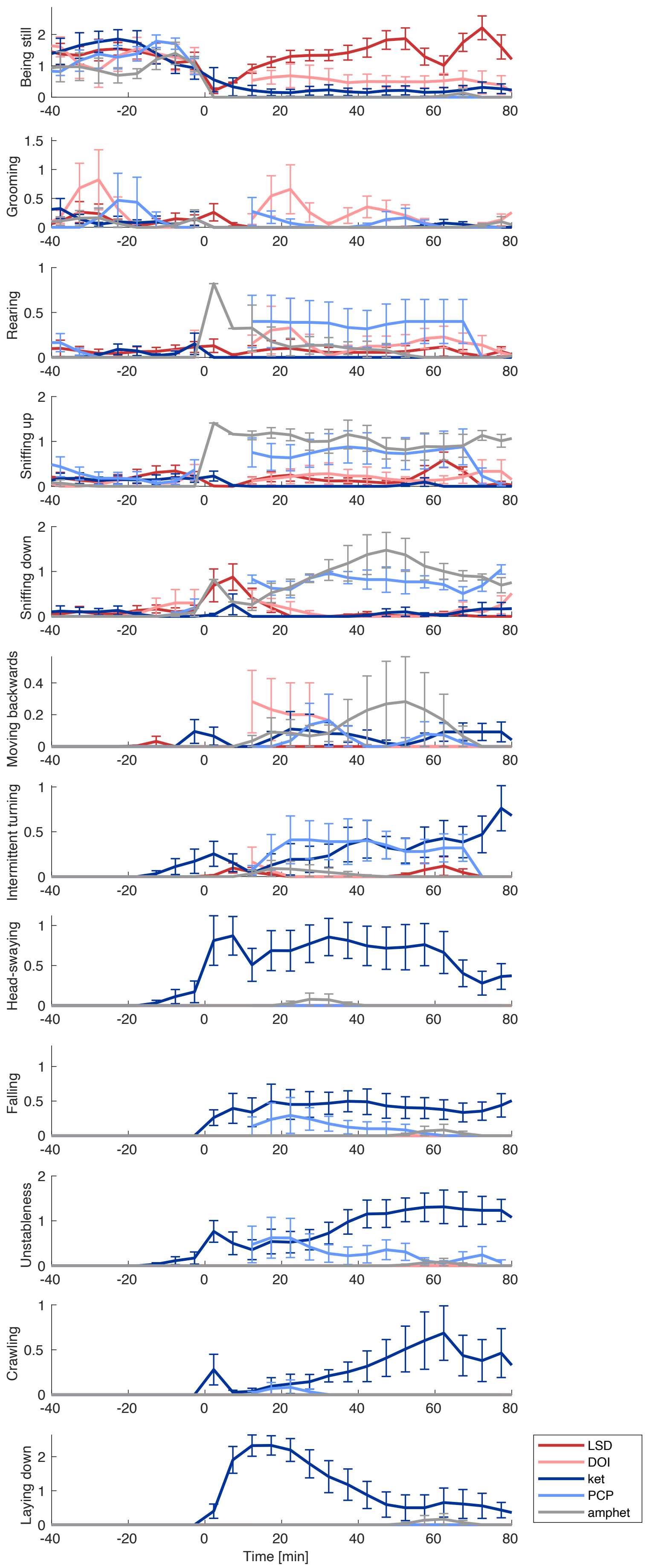


Time evolution of behaviors scored on a scale from 0 to 3 during 1 minute of observation each 10 minutes. Scores were interpolated in 5-minute steps and averaged across recording sessions. Lines show mean±SEM for LSD (red), DOI (pink), ketamine (dark blue), PCP (light blue) and amphetamine (grey). Times are relative to injection.

### Figure S2: Summary of manually scored behaviors

Average changes in behavior for each condition (Base = baseline, 2A = LSD or DOI, NMDA = ketamine or PCP, Am = amphetamine), scored on a scale from 0 to 3. Data was averaged over the periods [-35 -5] minutes for baseline and [30 60] minutes for the other conditions (relative to injection). Bars show mean and SEM, asterisks show significance at the p<0.05 (*), p<0.01 (**) and p<0.001 (***) levels (nested ANOVA).

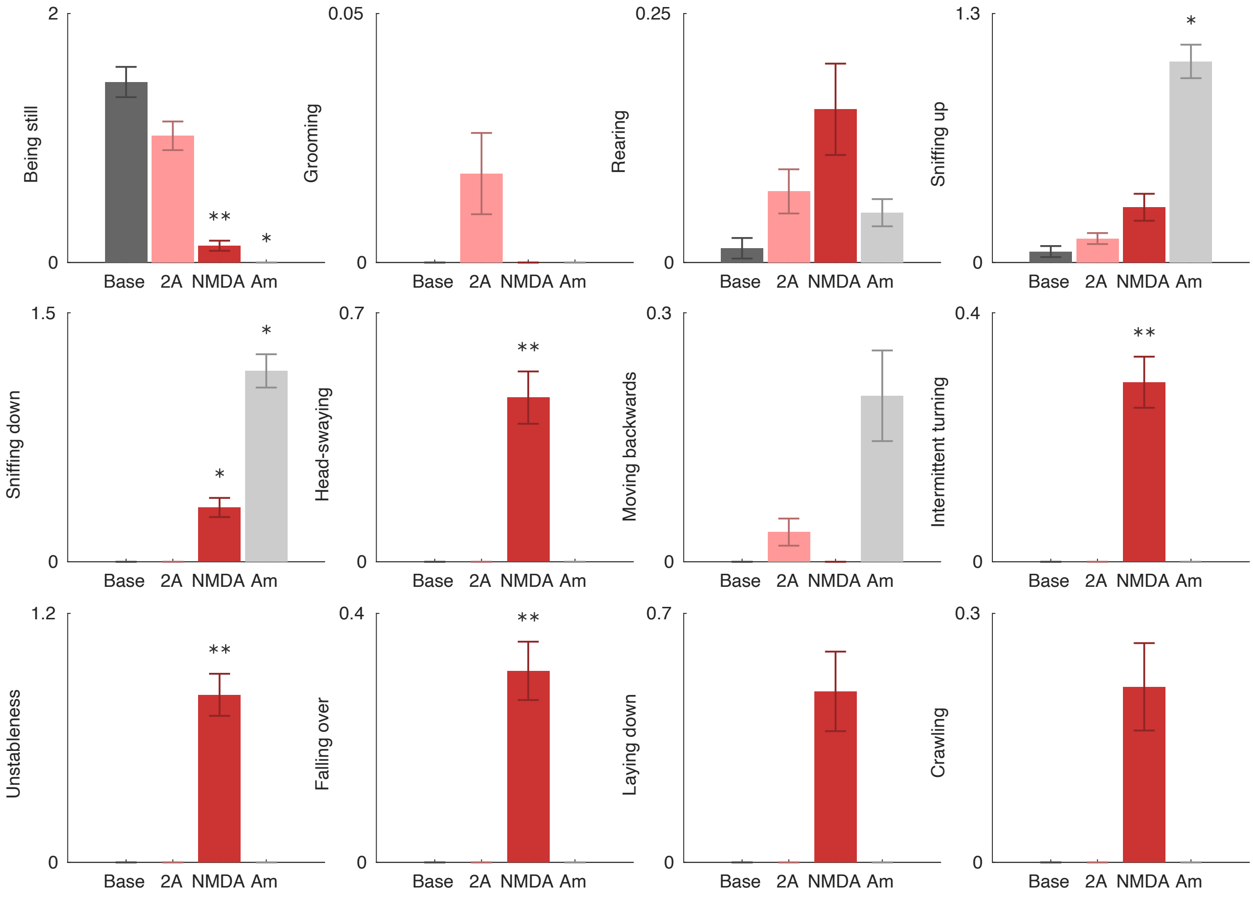

### Figure S3: Validation of head-twitch response detections

**A.** The medial-lateral (ML) component of the acceleration shows a clear oscillation during the HTR event.
**B.** The spectrogram of the ML component shows that the oscillation is found between 8 and 32 Hz (magenta dashed lines).

**C.** Absolute value of C smoothed with a Gaussian window ($\sigma$=50 ms).

**D.** Example data from an LSD recording showing high correspondence between manual and automatic detection. Manually identified events were divided in three classes based on their intensity: 1 = short side-to-side movement of the head but not a clear shake; 2 = clear shake of the head and/or part of the anterior portion of the trunk; 3 = clear powerful shake with most of the body involved. Colored diamonds correspond to manually identified HTR events (blue = class 1, not shown; purple = class 2; red = class 3). Grey diamonds indicate HTR events detected by the accelerometer with a threshold at 0.4 (grey dashed line).
**E.** Distribution of HTR indices for manually detected HTR events, split on manually identified HTR classes (grey = non-HTR; blue = HTR1; purple = HTR2; red = HTR3). There is a clear separation between HTR and non-HTR events (grey distribution), and a partial separation between HTR classes 1-3.
**F.** ROC curves with True Positives defined as manual scores >= 1 (blue line), manual scores >=2 (purple line) or manual scores = 3 (red line). The area under the curve (AUC) shows a near perfect classification for all three cases.

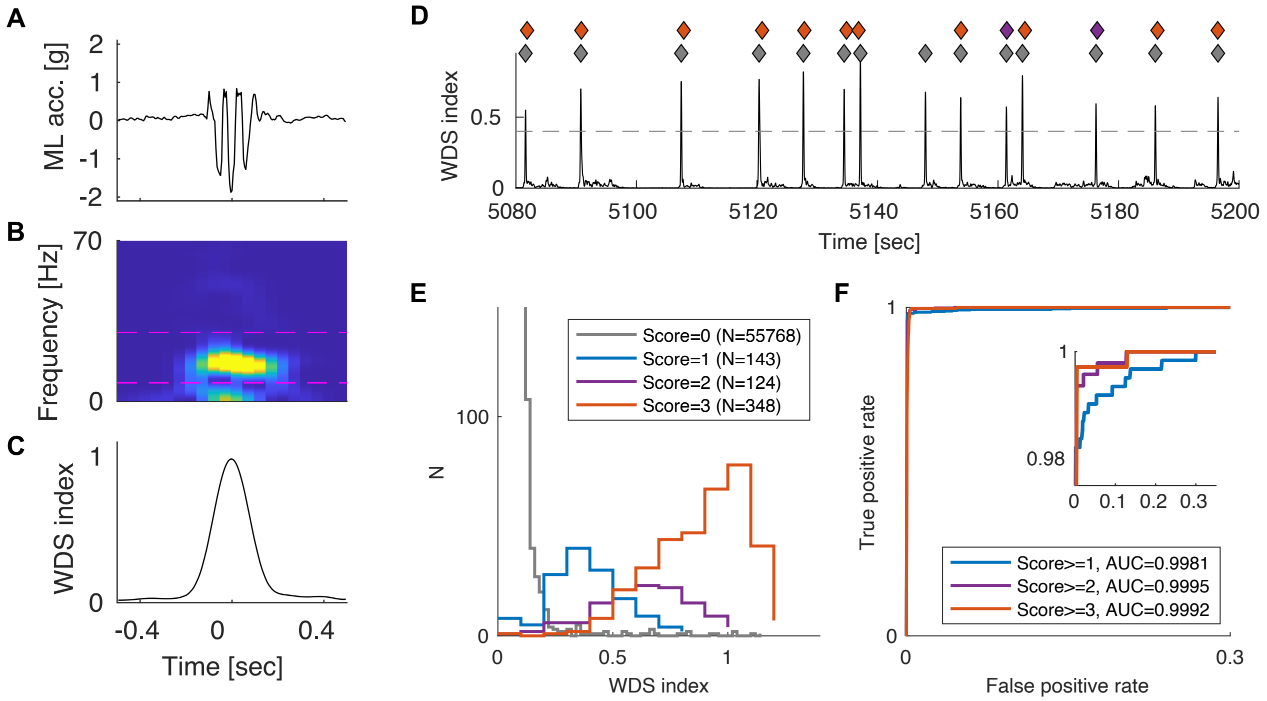

### Figure S4: Classification of units into putative cell types

Scatter plots of waveform features (left column) and corresponding waveforms (right column). Three features were used in the classification: peak width (FWHM), peak-to-valley time and valley width (FWHM; not shown). Yellow = putative principal cell, blue = putative interneuron, grey = unclassified.

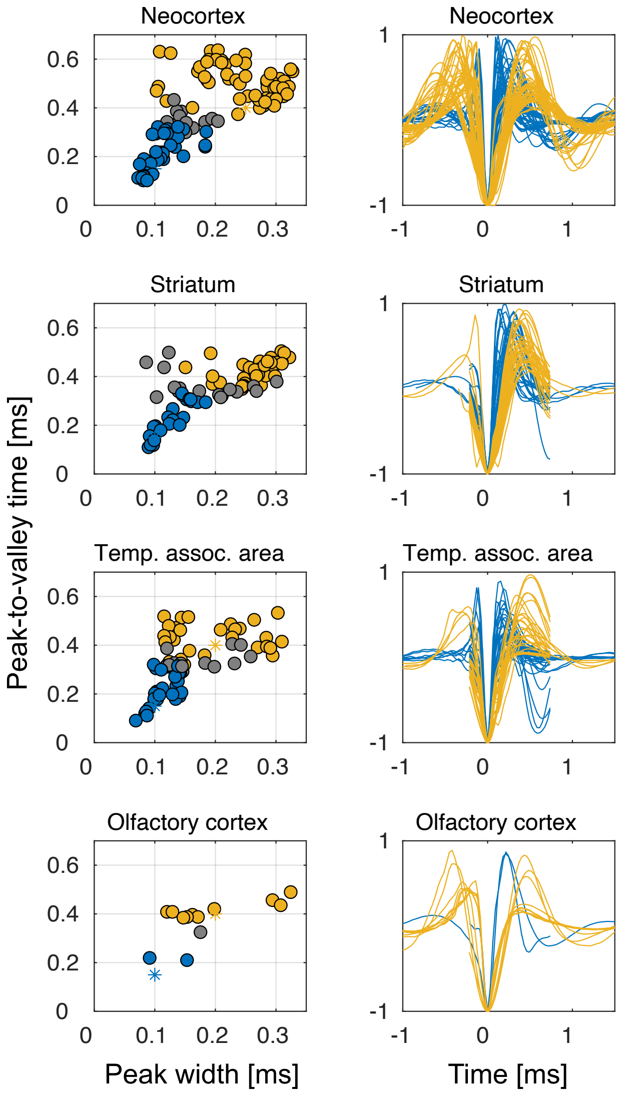

### Figure S5: Modulation of neuronal firing rates in response to amphetamine

**A.** Standardized neuronal firing rate responses to amphetamine. Each row shows the activity of a single unit and rows are rank ordered according to the response during the drug period (indicated by the black bar).

**B.** Average standardized neuronal firing rates (left panel) and fraction of modulated cells (right panel) in response to amphetamine for different cell populations (PC=putative principal cells, IN=putative interneurons, X=unclassified cells). Standardized rates for the 30 to 60 minutes post drug injection are compared to the -35 to -5 minutes baseline. In the right panel, the fractions of down-modulated cells are shown in red and up-modulated cells are shown in green. Asterisks indicate significance at the p<0.05 level (nested ANOVA) for rate modulations and p<0.05 (binomial test) for fractions. The numbers next to the cell labels indicate the number of cells in each population.

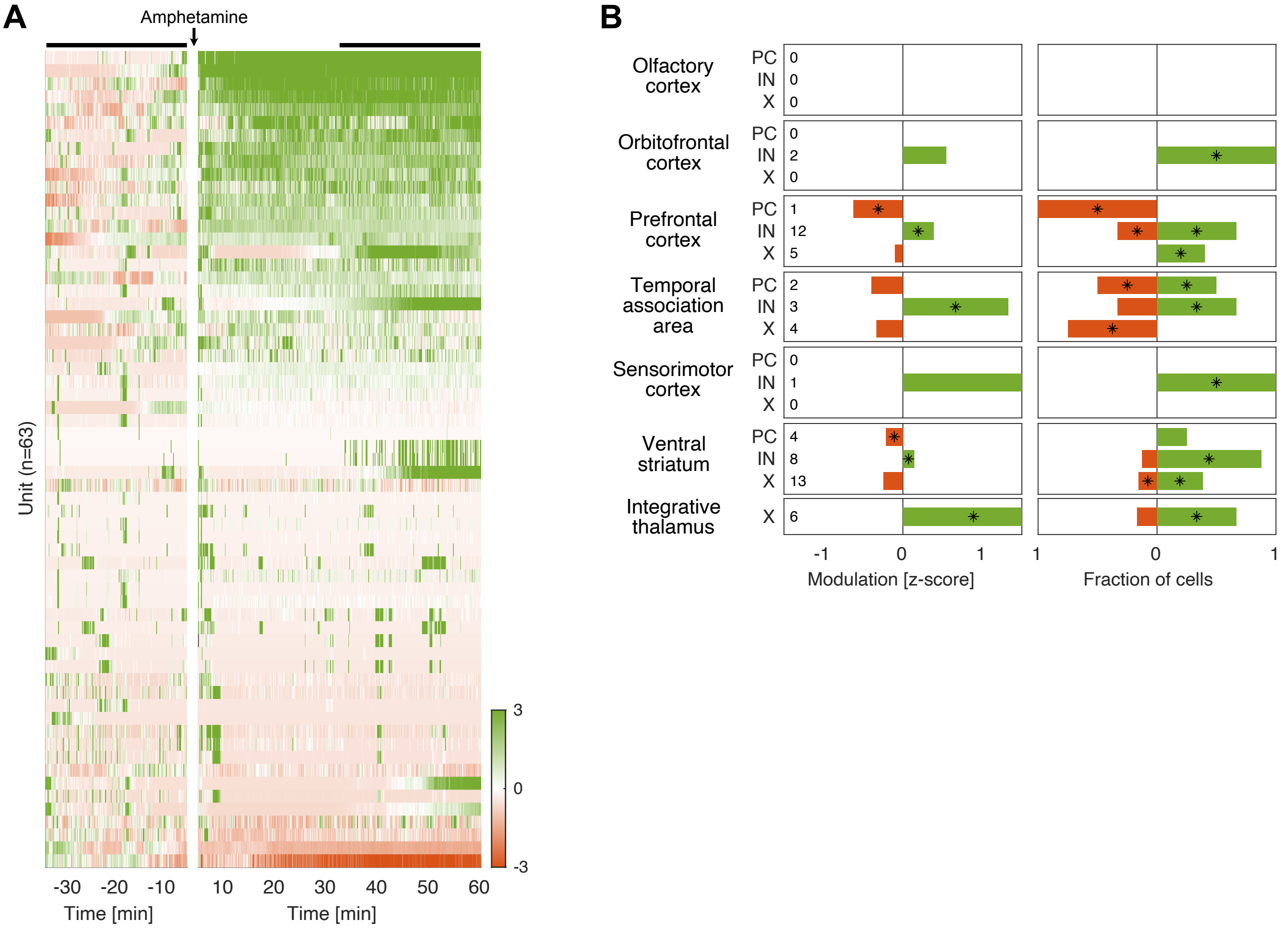

**Figure S6: Modulation of LFP spectra in response to amphetamine**

Power spectra of local field potentials during baseline (grey) and amphetamine (blue).

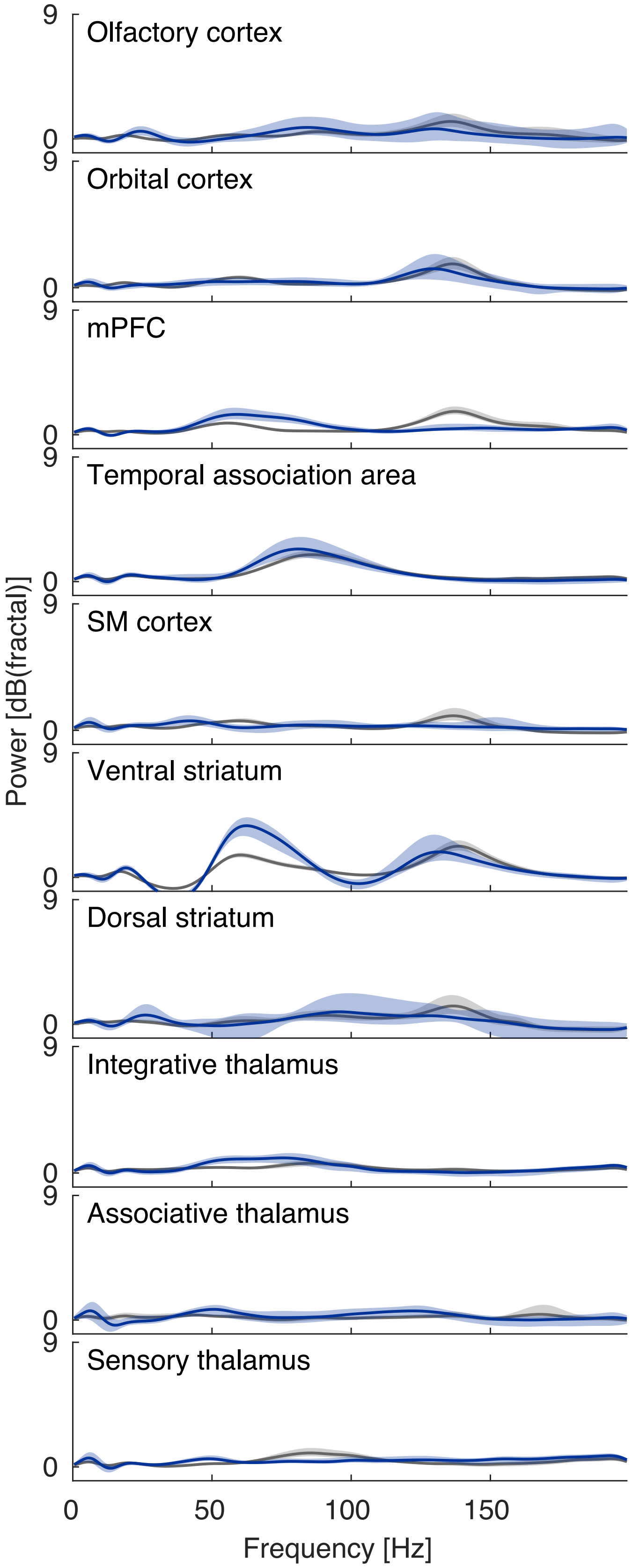

### Figure S7: HFO detection

**A.** Examples of LFP power spectra (grey) from 8 seconds of data. Red or green lines show the corresponding fitted function

$$y(f) = a_1 e^{\left(\frac{f-a_2}{a_3}\right)^2} + a_4 + a_5.$$

Functions with parameter values and goodness-of-fit within given limits were counted as successful HFO detections and marked in green.

**B.** Spectrograms (same as in Figure 3B) with successful HFO detections marked with green crosses.

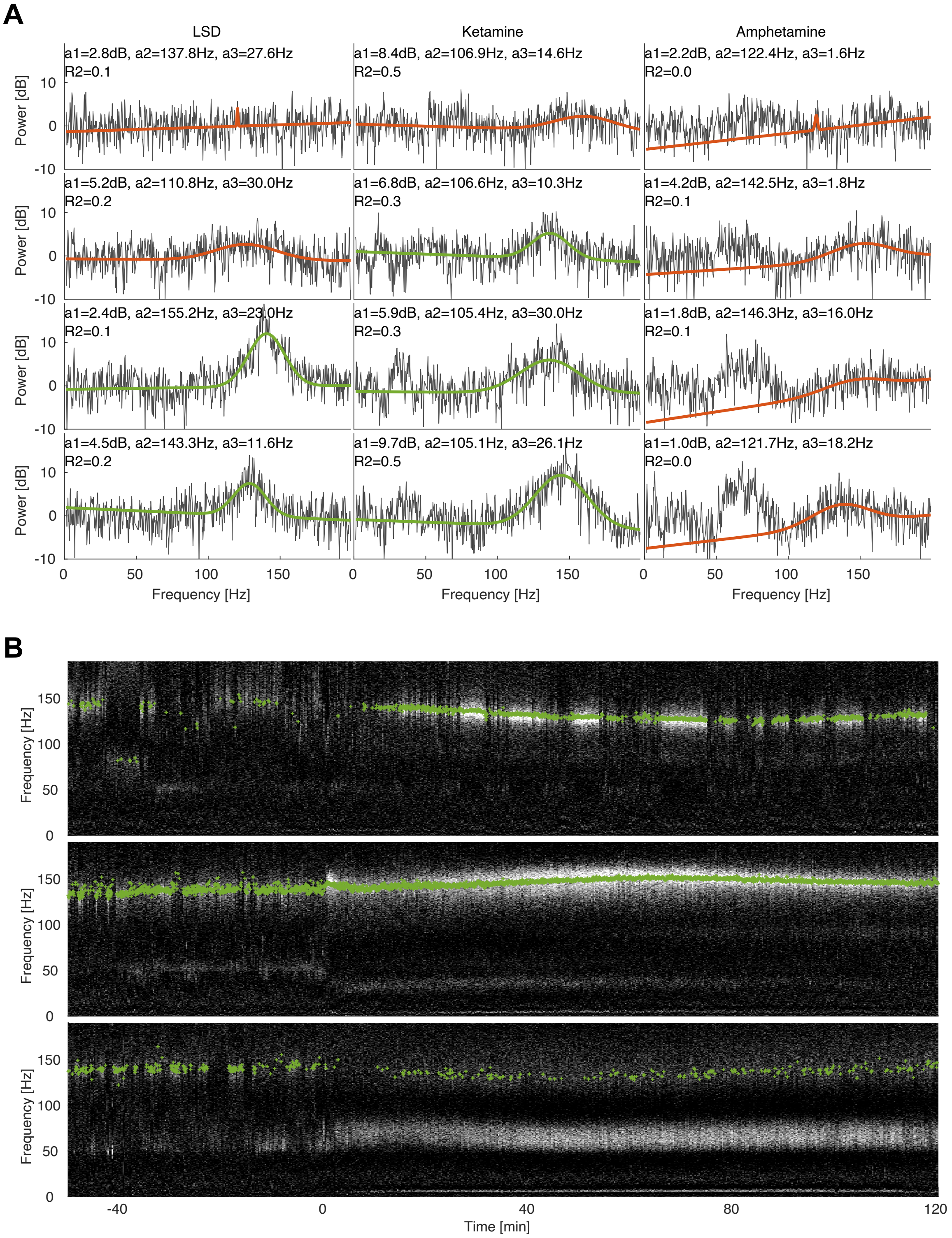

### Figure S8: Classification of HFO detection rates

To better visualize how prevalent HFOs were in different structures and recording sessions, we defined 4 prevalence classes based on the distribution of detection rates. First, the detection rate was determined for each recording session and treatment state. The fraction of sessions with a higher rate than a given threshold was then plotted as a function of that threshold (pink lines = 5HT2A, red lines = NMDA, dark grey lines = baseline, light grey lines = amphetamine). HFOs were classified as being Persistent if the detection rate was higher than 90% in at least 33% of sessions (red zone). They were Prevalent if the detection rate was higher than 50% in at least 33% of sessions (orange zone). Otherwise HFOs were Occasional (yellow zone), unless the detection rate was higher than 5% in no more than 5% of sessions. Then they were classified as Absent (grey zone).

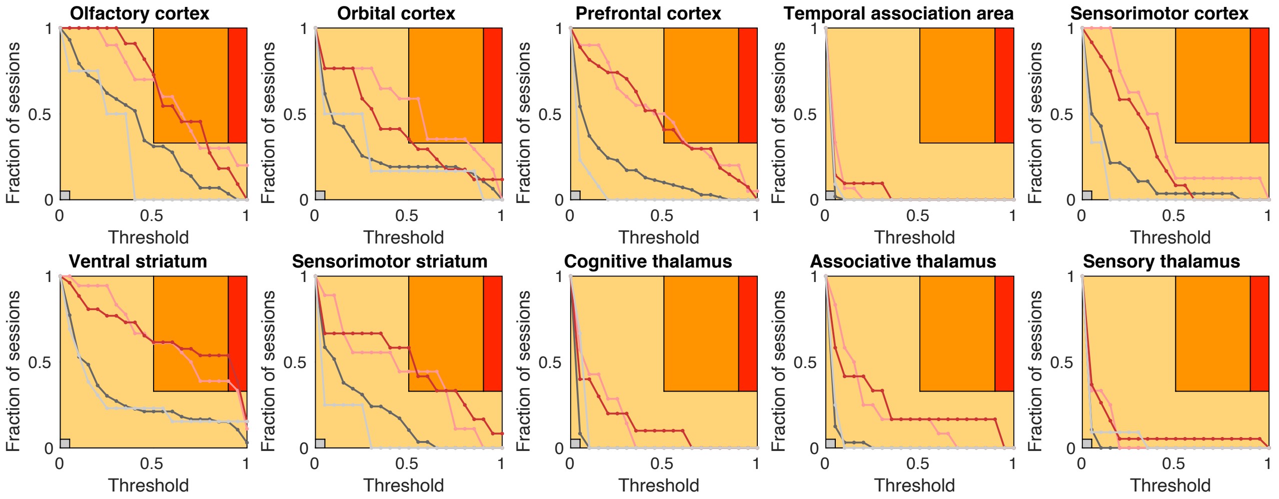
